# *GDCRNATools*: an R/Bioconductor package for integrative analysis of lncRNA, miRNA, and mRNA data in GDC

**DOI:** 10.1101/229799

**Authors:** Ruidong Li, Han Qu, Shibo Wang, Julong Wei, Le Zhang, Renyuan Ma, Jianming Lu, Jianguo Zhu, Wei-De Zhong, Zhenyu Jia

**Affiliations:** Department of Botany and Plant Sciences, University of California, Riverside, CA, USA; Genetics, Genomics, and Bioinformatics Program, University of California, Riverside, CA, USA; College of Animal Science and Technology, Nanjing Agricultural University, Nanjing, Jiangsu, China; Department of Mathematics, Bowdoin College, Brunswick, ME, USA; Department of Urology, Guangdong Key Laboratory of Clinical Molecular Medicine and Diagnostics, Guangzhou First People’s Hospital, the Second Affiliated Hospital of South China University of Technology, Guangzhou, China; Department of Urology, Guizhou Provincial People’s Hospital, Guizhou, China

## Abstract

The large-scale multidimensional omics data in the Genomic Data Commons (GDC) provides opportunities to investigate the crosstalk among different RNA species and their regulatory mechanisms in cancers. Easy-to-use bioinformatics pipelines are needed to facilitate such studies. We have developed a user-friendly R/Bioconductor package, named *GDCRNATools*, to facilitate downloading, organizing, and analyzing RNA data in GDC with an emphasis on deciphering the lncRNA-mRNA related competing endogenous RNAs (ceRNAs) regulatory network in cancers. Many widely used bioinformatics tools and databases are utilized in our package. Users can easily pack preferred downstream analysis pipelines or integrate their own pipelines into the workflow. Interactive *shiny* web apps built in *GDCRNATools* greatly improve visualization of results from the analysis.

**Availability:** *GDCRNATools* is an R/Bioconductor package that is freely available at https://github.com/Jialab-UCR/GDCRNATools

## 1. Introduction

Competing endogenous RNAs (ceRNAs) are RNA molecules that indirectly regulate other RNA transcripts by competing for the shared miRNAs. Deregulation of ceRNA networks may lead to human diseases, such as cancer (Gupta, et al., 2010; Ning, et al., 2015; Schmitt and Chang, 2016). Mounting evidences show that lncRNAs harboring multiple miRNA response elements (MREs) can act as ceRNAs to sequester miRNA activity and thus reduce the inhibition of miRNAs on their target genes (Kallen, et al., 2013; Salmena, et al., 2011). Although a few lncRNA-related ceRNAs have been reported to play critical roles in cancer development (Kumar, et al., 2014; Liu, et al., 2014), the regulatory mechanisms and significances of a large portion of ceRNAs remain to be unraveled.

The Genomic Data Commons (GDC) provides the cancer research community with a repository of standardized genomic and clinical data from National Cancer Institute (NCI) programs including The Cancer Genome Atlas (TCGA) and Therapeutically Applicable Research To Generate Effective Treatments (TARGET). High-quality datasets from non-NCI research programs such as genomic data from the Foundation Medicine company are also maintained in GDC. Advanced tools, such as *TCGA-Assembler* (Zhu, et al., 2014) and *TCGAbiolinks* (Colaprico, et al., 2016) that were initially developed for retrieving, processing and analyzing TCGA data from DCC (Data Coordinating Center) have been updated to access GDC data. However, none of these tools offer a route for a comprehensive analysis of RNA-seq and miRNA-seq data available in GDC.

Here we present a new R/Bioconductor package, named *GDCRNATools* to facilitate downloading, organizing, and integrative analyzing RNA data in the GDC (Fig. 1). Many analyses can be performed using *GDCRNATools* including differential gene expression analysis, ceRNAs regulatory network analysis, univariate survival analysis, and functional enrichment analysis. A newly developed algorithm *spongeScan* (Furió-Tarí, et al., 2016) is used to predict MREs in lncRNAs acting as ceRNAs. In addition, databases including *starBase v2.0* (Li, et al., 2013), *miRcode* (Jeggari, et al., 2012) and *miRTarBase 7.0* (Chou, et al., 2017) were also integrated and used as evidence basis for miRNA-mRNA and miRNA-lncRNA interactions. We updated gene IDs in these databases according to the latest Ensembl 90 annotation of human genome, and unified mature miRNA IDs based on the new release miRBase 21. *GDCRNATools* allows users easily perform the comprehensive analysis or integrate their own pipelines such as molecular subtype classification, weighted correlation network analysis (WGCNA) (Langfelder and Horvath, 2008), and TF-miRNA co-regulatory network analysis, etc. into the workflow.

**Fig. 1.**
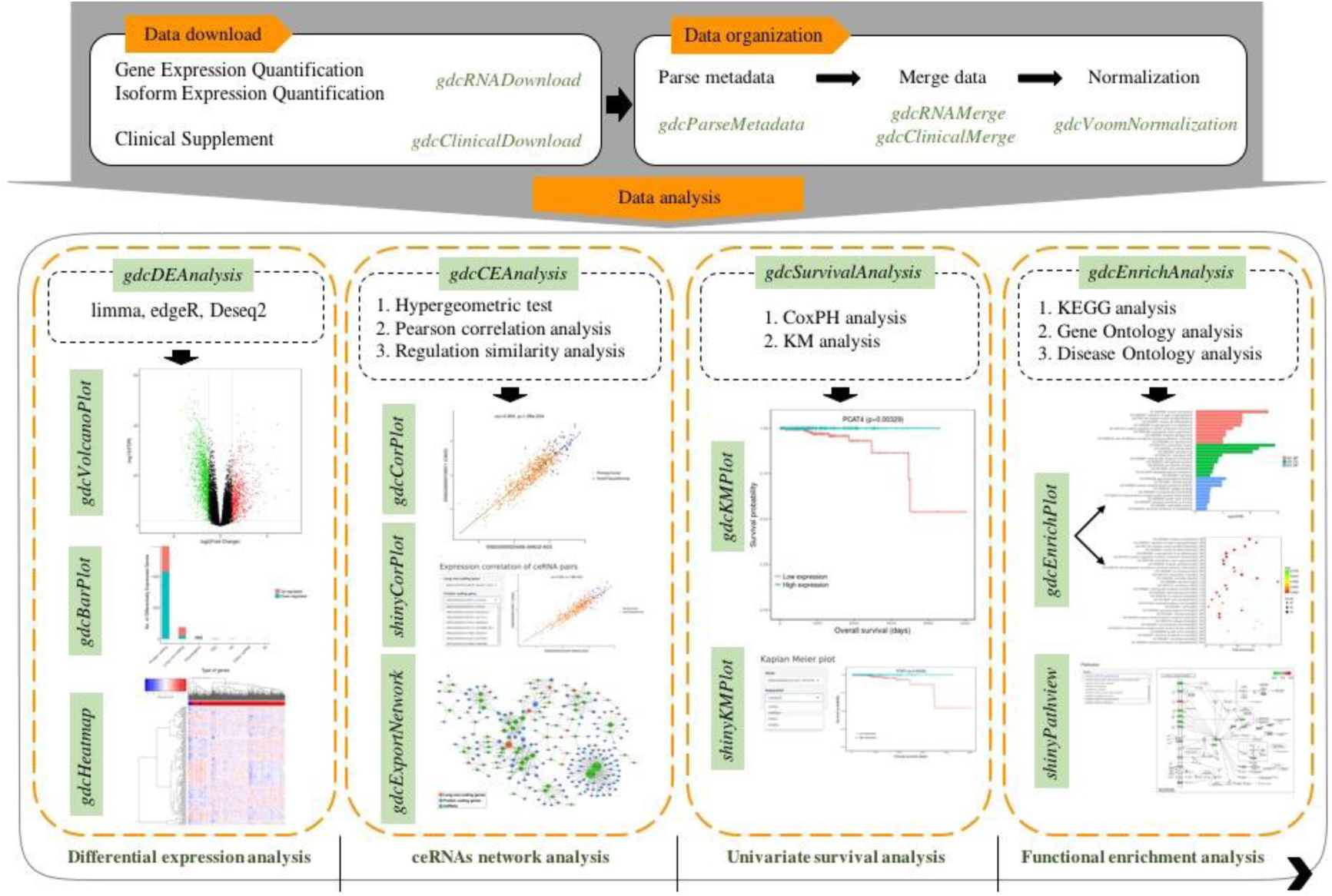
Workflow of GDCRNATools.

## 2. Implementation and main functions

### 2.1 Download and process data

Two strategies for downloading RNA expression data are available in the *gdcRNADownload* function: (1) users can simply provide the manifest file that contains Universally Unique Identifiers (UUIDs) corresponding to data in the GDC cart, or (2) download data automatically by specifying project id and data type. Metadata associated with downloaded files can be easily parsed by *gdcParseMetadata* to facilitate downstream analysis. *gdcRNAMerge* function merges total read counts for 5p and 3p strands of miRNAs (processed from isoform quantification data) and HTSeq read counts of gene quantification data into single expression matrix, respectively. Clinical data can be downloaded and processed by the *gdcClinicalDownload* and *gdcClinicalMerge* functions.

### 2.2 Differential gene expression analysis

Three most commonly used methods: *limma* (Ritchie, et al., 2015), *edgeR* (Robinson, et al., 2010), and *DESeq2* (Love, et al., 2014) can be implemented in *gdcDEAnalysis* function to identify differentially expressed genes (DEGs). Gene symbols and biotypes based on the Ensembl 90 annotation are reported in the output of *gdcDEReport*.

### 2.3 ceRNAs regulatory network analysis

Three criteria are used to define competing lncRNA-mRNA pairs: (1) the number of shared miRNAs by a lncRNA-mRNA pair and hypergeometric probability associated with this number, (2) the strength of positive expression correlation between lncRNA and mRNA, and (3) the overall regulation similarity of the lncRNA-mRNAs pair mediated by shared miRNAs, which is defined as:

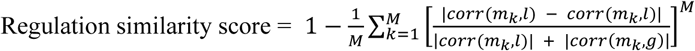

where *M* is the total number of shared miRNAs, *m_k_* is the *k*th shared miRNAs with *k* = 1, …, *M*, and *corr*(*m_k_,l*) and *corr*(*m_k_,g*) represents the Pearson’s correlation between the *k*th miRNA with lncRNA, and with mRNA, respectively. Three miRNA-mRNA interaction databases (*StarBase v2.0* (Li, et al., 2013), *miRcode* (Jeggari, et al., 2012), *mirTarBase 7.0* (Chou, et al., 2017)) and three miRNA-lncRNA interaction databases (*StarBase v2.0* (Li, et al., 2013), *miRcode* (Jeggari, et al., 2012), *spongeScan* (Furió-Tarí, et al., 2016)) are incorporated and used in the *gdcCEAnalysis* function internally. *gdcCEAnalysis* also allows user-provided data of miRNA-lncRNA or miRNA-mRNA interactions (either predicted using other algorithms or validated through experiments) to be utilized for the ceRNAs regulatory network analysis. The resultant lncRNA-miRNA-mRNA interaction network can be exported by the *gdcExportNetwork* function and then visualized in *Cytoscape* (Shannon, et al., 2003).

### 2.4 Functional enrichment analysis

*gdcEnrichAnalysis* performs Gene Ontology (GO) and Kyoto Encyclopedia of Genes and Genomes (KEGG) functional enrichment analyses using the latest databases through the R/Bioconductor package *clusterProfiler* (Yu, et al., 2012). Disease Ontology analysis using *DOSE* package (Yu, et al., 2014) is also included in the *gdcEnrichAnalysis* function to detect gene-disease associations.

### 2.5 Survival analysis

Two methods are available in *gdcSurvivalAnalysis* function to perform univariate survival analysis: (1) Cox Proportional-Hazards (CoxPH) regression and (2) Kaplan Meier (KM) analysis. *gdcSurvivalAnalysis* takes a list of genes as input and reports the hazard ratio, 95% confidence intervals, and p value of significance test on overall survival of each gene.

### 2.6 Visualization

In addition to the routine visualization methods, such as volcano plot, heatmap, KM plot, etc., a more attractive visualization feature of *GDCRNATools* is the application of interactive *shiny* web apps, which allow users to view survival curves, expression correlation between lncRNA and mRNA, and enriched pathways by simply selecting genes/pathways of interests on the local webpage.

## 3. Conclusion

We have developed a novel R/Bioconductor package, named *GDCRNATools* to conduct advanced analyses of RNA-seq and miRNA-seq data in GDC data portal for identification of lncRNA-miRNA-mRNA competing triplets in cancer. This easy-to-use package allows users with little coding experience to perform the entire analyses smoothly. As standardized data from other programs may be submitted to the GDC, we believe that *GDCRNATools* will gain ground in cancer research for deciphering the crosstalk among different RNA species and their regulatory mechanisms.

